# MICRO-TAG enzyme complementation enables quantification of cellular drug-target engagement in temperature series

**DOI:** 10.1101/2025.09.04.674281

**Authors:** Ivan Babic, Nikolas Bryan, Claire Cunningham, Avery Sampson, Daniel Starczynowski, Elmar Nurmemmedov

**Affiliations:** CellarisBio, San Diego, CA, 92121, United States; Division of Experimental Hematology and Cancer Biology, Cincinnati Children’s Hospital, Cincinnati, OH, 45229, United States; Department of Cancer Biology, University of Cincinnati, Cincinnati, OH, 45267, United States; Department of Pediatrics, University of Cincinnati, Cincinnati, OH, 45229, United States; University of Cincinnati Cancer Center, Cincinnati, OH, 45267, United States

**Keywords:** Drug, cellular target engagement, temperature series, real-time

## Abstract

Drug discovery for challenging drug targets necessitates the proteomic complexities of the cellular milieu for contextual target folding and function. Conventional biophysical methods for assessing drug interaction with a target are often not sufficiently suited for drug discovery as they impose acellular environment on the target and rely on recombinant purified protein material. In contrast, cell target engagement offers a powerful paradigm for drug discovery, through measurement of transitions in the thermodynamic state of a target protein, as it engages with drug molecules in the cell. Split-enzyme cell target engagement methods offer scaled utility during early drug discovery. Here, we describe a novel highly sensitive and scalable fluorescence-based cell target engagement method that leverages complementation of split-RNase S. This offers a unique combination of procedural and biophysical advantages, enabling its seamless integration with various instruments and applications designed for fluorescence detection. Most importantly, this new method allows for quantitation of cell target engagement in programmable temperature series format, consistent with conventional thermal shift assays, rather than at a single melting temperature. We demonstrate the sensitivity and versatility of this approach for drug discovery using targets MAPK1, KRAS, and UBE2N.

## INTRODUCTION

Demonstrating direct engagement of a drug with its respective target is a key step in drug discovery. Advent of biophysical methods such as surface plasmon resonance (SPR), micro-scale thermophoresis (MST) and differential scanning fluorimetry (DSF) has enabled reliable quantitation of target engagement of drug candidates with isolated proteins under controlled conditions [1, 2]. Thermal shift has been widely adopted as a biophysical method to assess drug-induced conformational stabilization of proteins as they undergo a temperature-induced transition from a native folded to a denatured unfolded state. This is measured by a shift in the melting temperature of a protein (T_m_), which is defined as the temperature at which half the population is melted/denatured. Dye-based DSF method uses thermal scanning to monitor stability of purified recombinant proteins, on real-time PCR instruments [3].

Most drug targets, however, are not amenable to standard biophysical methods of target engagement, due to the artificial acellular environment employed by these methods. Such targets require the complexities of the physiological cellular environment for contextual folding and function. Drug-target binding activity in a cellular environment can be attributed to many factors including cell permeability, off-target binding, compound efflux, or a multiple structural and functional states of the target in the cell [4–6]. Cell target engagement (CTE) has emerged as a powerful paradigm in drug discovery for assessing the direct engagement of a drug with its target in a physiological context. CTE measures shifts in the thermodynamic state of the target protein and the induced biophysical events (stabilization and de-stabilization), as it interacts with drug molecules in a complex environment of the cell [7, 8]. CTE offers a significant advancement over DSF, as it enables measurement of drug-induced target stability within the native milieu of the cell. CTE allows measurement of drug-target engagement by quantifying the amount of folded protein remaining in the cell in response to a heat stress, typically at the temperature of aggregation, T_agg_50, a point at which half the target protein population exists under aggregated stage. CTE measures the shift in biophysical stability of a target protein induced by a drug molecule [9].

Several split-enzyme CTE applications have recently emerged to offer sensitive and scalable, quantitation for drug-target engagement [10, 11]. These strategies involve enzyme complementation systems whereby a peptide tag, usually a small subunit of larger enzyme, is cloned into the N- or C-terminus of the target protein of interest. The nature of the cloned tag is selected to readily complement with a large subunit, which is added exogenously as a free reagent. A holoenzyme is formed upon complementation within in the cell, and the processing of added substrate generates a detectable signal proportional to the level of available target protein. Application of temperature challenge to the cells induces aggregation of the target protein, which can be rescued upon the direct engagement of a drug molecule. Based on this principle, cellular target engagement potency and selectivity of therapeutic molecules can be measured [12, 13]. In the case of the HiBiT system, enzyme complementation forms a functional luciferase enzyme that allows quantification of soluble protein using luminescence for scaled testing of drug candidates [11, 14].

The currently available luminescence-based CTE methods, while highly valuable, are limited by operational hurdles, which includes multi-step sample processing, thereby compromising scalability and accountability desired for fast drug discovery workflows [15]. Conventional CTE strategies also rely on the exact determination of T_agg_50 point for target protein, followed by interrogation of ligand binding at that specific temperature after a prolonged incubation. CTE measurement under a defined T_agg_50 point provides a static and rather simplified view of thermodynamic profile of drug targets. Furthermore, it introduces potential bias for calculation of compound potency, as well as adds an extra step to the workflow, thus limiting system automatability. Traditional protein-based thermal shift assays employ temperature scanning to monitor drug binding and current CTE methods interrogate binding only at one temperature. T_agg_50 values can be highly dependent on the specific experimental conditions, target expression levels, numbers and health status of the used cells and precision of temperature ramping, potentially leading to variations in the T_agg_50 calculations [16, 17]. These limitations narrow its utility for certain biological applications and integration with readily available lab instruments. To overcome these hurdles, it is advantageous to remove the requirement for precise determination of the T_agg_50 value and to reduce the number of processing steps. Stepwise controlled aggregation of target proteins under temperature gradient can expose various conformational states, potentially imitating the plurality of the structural transitions they undertake. Temperature scanning, combined with the advantage of CTE methods, offer a novel method to potentially capture multi-state drug-target engagement events, and thus further advance drug discovery and development practices.

The MICRO-TAG system combines the benefits of conventional DSF and CTE methods, by enabling monitoring of cellular protein stability under temperature series. Proteins exist in the cell under various structural states reflecting their context-dependent functional assignments. Under such structural transitions, a protein might exhibit varying binding affinities for the same drug molecule. Conversely, target proteins with multiple ligand binding sites may exhibit them in conformation-dependent manner, thus further emphasizing the importance of dynamic structural transitions. Under such multivalent target engagement scenarios, the overall interaction can involve multiple, distinct binding events occurring with different affinities. Analysis of the individual phases in the binding curve can shed light on the involvement of specific domains or sites and their contribution to the overall binding strength. Capturing multi-state events in CTE setting can reveal such distinct drug-target engagement events, each corresponding to a specific protein conformation interacting with the drug molecule [18, 19]. Integration of the MICRO-TAG method with the thermal programmability of highly sensitive real-time systems eliminates the necessity for T_agg_50 determination prior to testing ligand binding. Assessment of ligand binding in a temperature series offers convenience by decreasing the number of processing steps involved in setting up the reaction.

Based on a novel fluorescence-based CTE method, the MICRO-TAG system is built on complementation of split-RNase S. The MICRO-TAG enzyme complementation system has already been adopted by several recent drug discovery studies [20–22]. The method enables interrogation of drug targets without observable interference with their thermal melting profile – all within the physiological milieu of the cell. Importantly, this new method allows for quantitation and monitoring of cell target engagement using real-time instruments programmed to monitor protein levels in controllable temperature series format. The MICRO-TAG methods can be readily integrated with widely available real-time instruments, thus making it highly sensitive, programmable and scalable with fewer procedural steps. This novel method captures drug-target engagement events at various thermodynamic states, hence potentially enabling measurement of multi-state target engagement events and offering applicability for structurally diverse challenging drug targets. We demonstrate the utility of this strategy for drug discovery with data for protein targets MAPK1, UBE2N and KRAS.

## RESULTS

### Assembly of MICRO-TAG system for cell target engagement

MICRO-TAG cell target engagement method improves on the previously reported enzyme complementation methods by integrating an enzymatic reaction for fluorescence development. Assembly of a reporter cell line starts with cloning of a 15-amino acid tag to either N- or C-terminus of a target protein of interest. With favorable biophysical properties (**Table S1**), this tag corresponds to the S-tag peptide (KETAAAKFERQHMDS) derived from bovine pancreatic ribonuclease S (RNase S) [23]. The S-tag is the small subunit of a two-subunit ribonuclease S (RNase S). RNase S is a complex that consists of two proteolytic fragments of bovine pancreatic RNase S: the S-peptide (residues 1-20) and S-protein (residues 21-124). The minimal sequence of the S peptide for enzyme complementation with the large S protein is amino acids 2 to 16 [24]. While S-tag is directly cloned to the target protein of interest, recombinantly expressed and purified S-protein is provided exogenously for signal detection (**Fig S1 and S2**). Complementation of S-tag with the large S-protein subunit forms an active RNase S protein capable of cleaving a specific RNA sequence (**Fig 1A and 1B**). Active RNase S enzyme cleaves an RNA sequence specific FRET substrate to generate fluorescence signal. The FRET substrate is an optimized fluorogenic substrate, 6-FAM-dArUdAdA-6-TAMRA, where 6-FAM refers to 6-carboxyfluorescein fluorophore and 6-TAMRA refers to 6-carboxy-tetramethylrhodamine fluorophore (**Fig 1B**). The option to select donor-quencher FRET pair fluorophores flanking this specific RNA sequence provides a highly sensitive and versatile strategy for quantifying enzyme complemented RNase S (**Fig 1B**).

**Figure 1:**
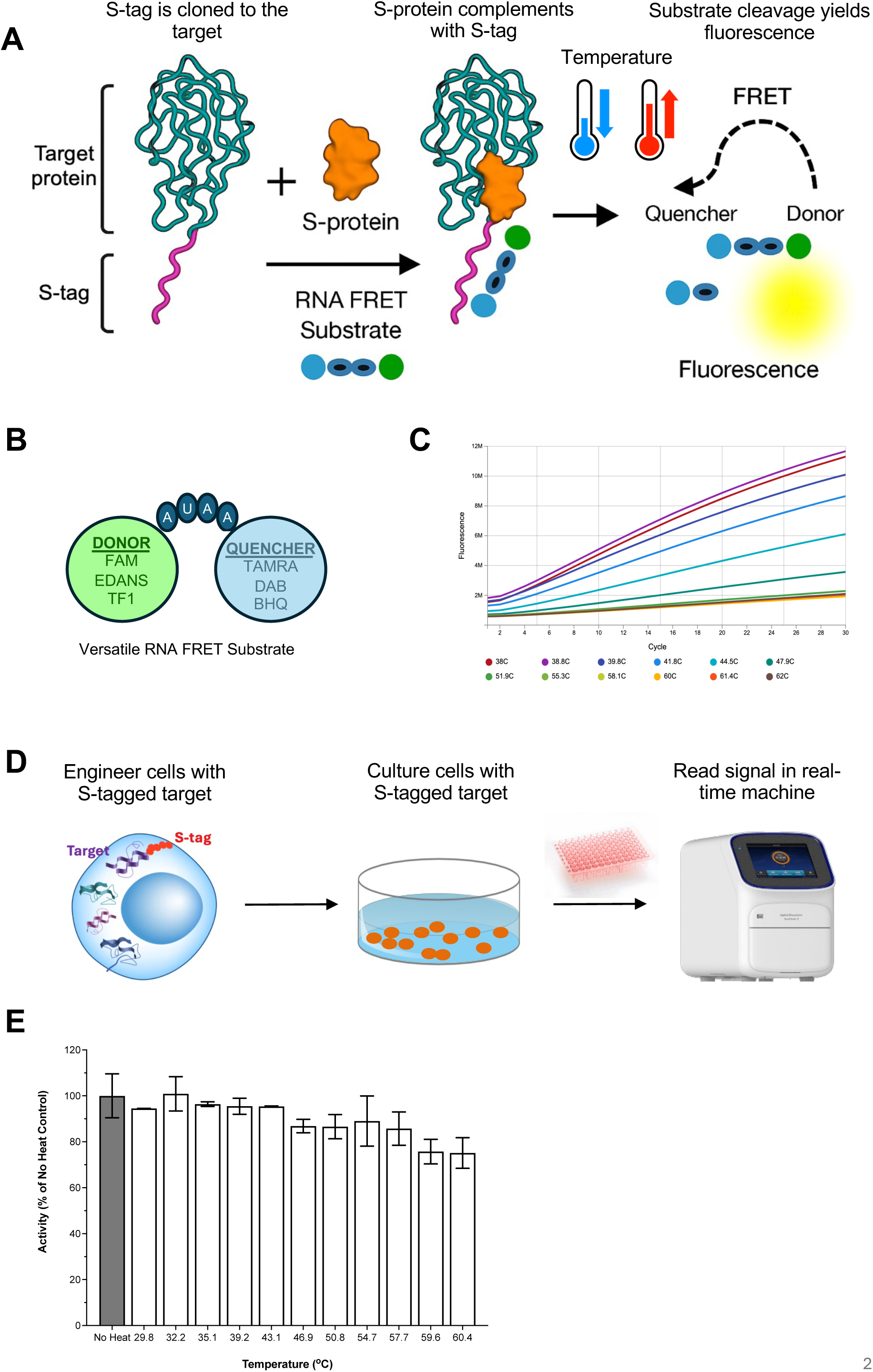
Assembly of MICRO-TAG real-time cell target engagement system. (A) Schematic representation of the MICRO-TAG enzyme complementation system. Target protein is shown in green, S-tag in purple, S-protein in orange, FRET RNA substrate as line of circles, and temperature control with blue and red arrows. (B) Schematic representation of versatility of the FRET RNA substrate. Various options for donors and quenchers are depicted inside large circles. (C) Representative real-time fluorescence curves from qPCR real-time system quantifying soluble MTH1-S-tag fusion when cells are exposed to various temperatures. (D) Schematic of the streamlined workflow for temperature series cell target engagement, showing the process from engineering and culturing the MICRO-TAG reporter cells to signal measurement in real-time instrument. (E) Thermal stability of recombinant S-protein. Enzymatic activity of S-protein at each tested temperature is shown in comparison to no-heat control. The experiment was performed in triplicate and error bars represent standard deviation of the mean.

The MICRO-TAG enzyme complementation system is designed as a high-throughput method for sensitive quantitation of unaggregated protein remaining after a heat challenge. **Fig 1C** demonstrates the sensitivity of this method in a qPCR real-time system where MTH1-S-tag-expressing cells are subjected to escalating temperature challenge followed by addition of the S-protein and FRET RNA substrate to quantify the amount of unaggregated MTH1 fraction. The temperature-dependent signal reduction shows significant decrease in fluorescence with increasing temperature, thus demonstrating the utility and sensitivity of this system in real-time instrument setting.

Combination of the MICRO-TAG system with real-time instruments streamlines cell target engagement (CTE) workflows by removing procedural complexity (**Fig 1D**). This new system takes advantage of precision programmability of real-time instruments to enable measurement of cell target engagement in temperature series. The method eliminates the conventional requirement to determine the T_agg_50 point of the target protein prior to ligand testing. This is made possible by systematically scanning various temperature points against multiple doses of the ligand. Programmed temperature series together with the “mix-and-detect” feature of the MICRO-TAG method offers convenience by decreasing the number of procedural burden and temperature bias imposed by the conventional CTE assays. Sequential escalation of temperature is possible owing to high thermal stability of RNase S in heat challenge, combined with temperature gradient programmability of real-time instruments. To demonstrate stability of the S-protein, we subjected the detection reagents (S-protein and FRET RNA substrate) to heat challenge of 30°C-60°C prior to adding to the MTH1-S-tag reporter cells. This revealed high thermal stability of S-protein with only 10% drop of enzymatic activity at the highest temperature points (**Fig 1E**).

### Workflow of MICRO-TAG cell target engagement temperature series

Use of the MICRO-TAG system with real-time instruments to monitor drug binding in a temperature series, which more closely resembles conventional thermal shift assays, is enabled by the heating programmability of real-time systems. In the MICRO-TAG workflow, cells expressing the target are combined with detection reagents in a PCR plate in the presence of 0.25% Triton-X-100 to allow the reagents to enter the cells. The real-time instrument is programmed as shown in **Fig. 2A**. During the temperature ramping step, a desired T (temperature) point is reached at a pre-determined ramping speed (e.g. 1.6 °C /sec). Once the temperature point is reached, the sample is incubated for a desired period (e.g. 3 minutes). During this step, the target will undergo thermodynamic shift under the thermal pressure. The temperature is ramped back to 25°C, at which the drug-target pair will equilibrate under the newly imparted structural perturbation and fluorescence signal from cleavage of FRET substrate is detected. The temperature is lowered to 25°C after heating to allow FRET RNA substrate cleavage since the split-RNase S enzyme is not active at higher temperatures. The ramp-incubate-detect cycle will repeat in a step-gradient fashion until the desired highest temperature point is reached.

**Figure 2:**
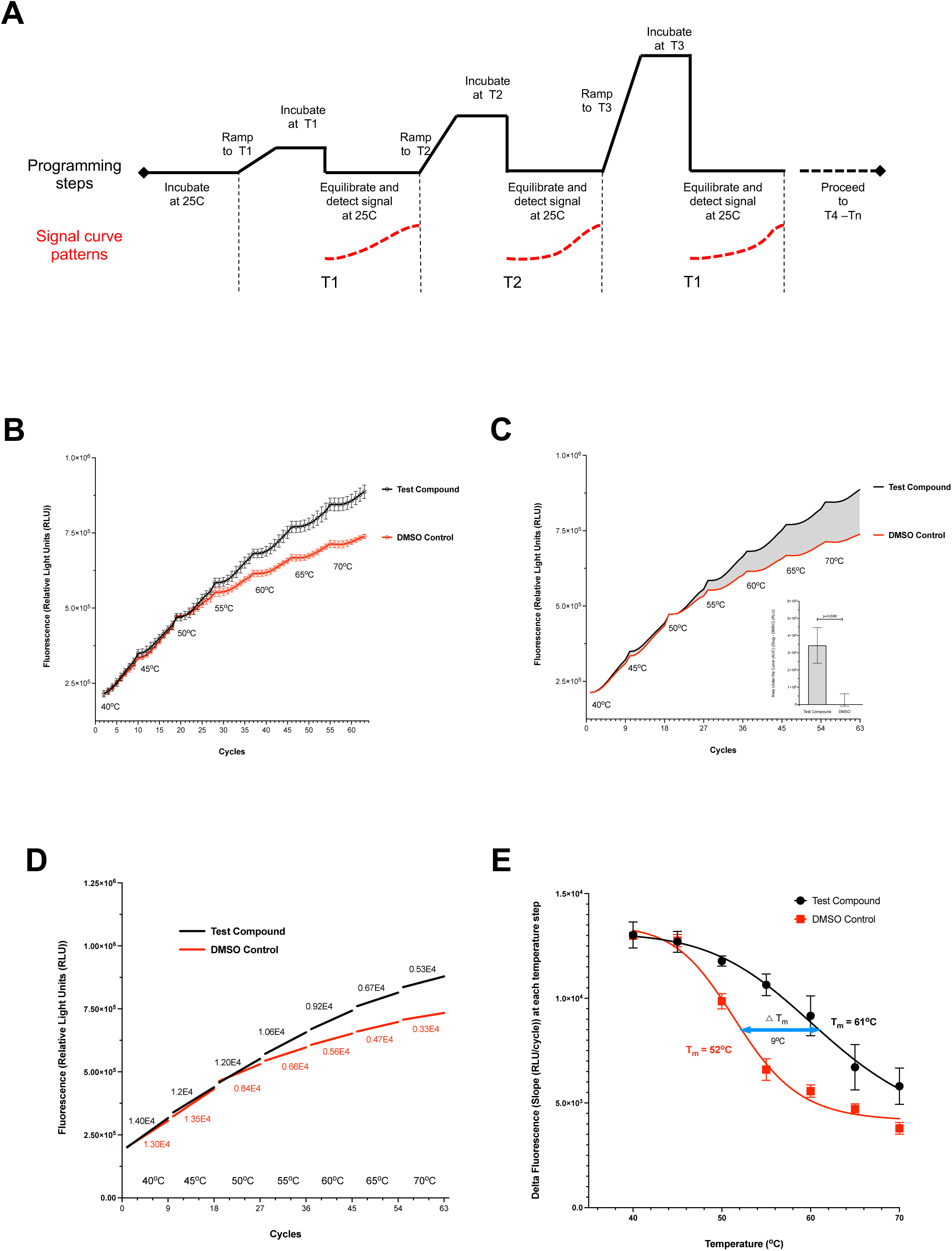
Workflow and output of MICRO-TAG real-time temperature series. (A) Programming of real-time qPCR instrument for use with MICRO-TAG in temperature series. The diagram shows the ramp-incubate-detect cycles together with expected signal patterns. (B) Representative readout from real-time temperature series system measured S-Crizotinib (test compound) binding and stabilizing MTH1. Temperature series profile of DMSO control is plotted together with S-Crizotinib. The experiment was performed in duplicate and error bars represent standard deviation of the mean. (C) Area Under the Curve (AUC) calculation for target stabilization by S-Crizotinib (test compound) relative to DMSO from the curves in panel B. The inset histogram is a quantification of AUC for both test compound and DMSO minus mean of the DMSO control. (D) Linear regression analysis of each detection step at specific temperatures from (B). Slopes are indicated above each temperature point. (E) Delta fluorescence analysis of data from (B). T_agg_50 values extracted from each curve together with delta-T are indicated.

Every tested temperature point imparts a gradual structural perturbation onto the target, thus stressing its engagement with the ligand. Signal detection at 25°C after every temperature challenge ensures that equilibrium is re-established, also providing consistency for signal development. The method enables CTE measurement using programmable temperature series rather than testing at a single temperature. As an example of the utility of the system for cell target engagement, the MICRO-TAG reporter cells expressing MTH1-S-Tag fusion were dispensed in 96-well plate at predetermined density. Drug molecule (S-Crizotinib) was added to the plate at desired final doses together with FRET substrate and S-protein and transferred to a real-time qPCR instrument. The MTH1-S-Tag fusion subjected to various temperatures displayed various conformational states, which, in turn, translates to corresponding levels of target abundance and hence FRET cleavage (**Fig 2B**). Calculation of the area under the curve (AUC) can be performed to estimate the magnitude of target stability induced by the drug at the tested dose (**Fig 2C**). At temperatures between 40°C-50°C, both DMSO control and the test compound S-Crizotinib (a well-characterized binding ligand of MTH1) followed an identical kinetic slope of 1.2-1.4×10^4^ RLU/cycle determined from linear regression of the detection steps (**Fig 2D**) [25]. At temperatures above 50°C, the signal in presence of the test compound (slope of 1.2×10^4^ RLU/cycle), continued to generate significant fluorescence signal, while DMSO control treated cells (slope of 0.8×10^4^ RLU/cycle), displayed less fluorescence. Signal increase for the DMSO control halved at 55°C, while that of the positive control S-Crizotinib persisted until 60°C. At the highest temperature point of 70°C, signal from both treatments plateaued, ultimately leaving a large area between the two curves, which indicates the magnitude of target stabilization by the drug. Plotting of the slopes (RLU/cycle) for each temperature point can be used to identify the temperatures impacting the thermal stability of the target (**Fig 2E**). In addition, it can be used to demonstrate the shift in melting temperature (Tm) of the target, which in the case of MTH1 corresponded to 52°C and 61°C, for DMSO and S-Crizotinib, respectively (**Fig 2E**).

To summarize, MICRO-TAG temperature-series data can be analyzed using either an AUC-based or a slope-based approach (see Methods). Both yield dose–response curves from which one can derive per-temperature EC50 values and an aggregated EC50_total_. Per-temperature EC50 is the dose that yields 50% of the modeled effect within a single temperature segment, while EC50_total_ is the dose that yields 50% of the aggregated effect across multiple temperature segments.

### MAPK1 case study: Cell Target Engagement Potency of AZD0364

MAPK1 / ERK2 is a challenging drug target because of its crucial role in many cellular processes, its high degree of structural similarity to ERK1, complex feedback loops within the MAPK signaling pathway that can lead to resistance mechanisms, and the potential for severe side effects due to its involvement in normal physiological functions across various tissues [26]. Here, we engineered MICRO-TAG reporter cells for MAPK1 by transient expression of human MAPK1 (Uniprot: P28482) with S-tag fusion in HEK293 cells. The expression of this engineered construct resulted in a single band of 41 kDa on an S-Tag immunoblot (**Fig 3A and Fig S3A**). Subsequent MICRO-TAG enzyme complementation assay yielded signal of around 8×10^5^ fluorescence units over the untransfected cells (**Fig 3B**). Additional verification of target expression and localization was confirmed by live-cell imaging using a modified MICRO-TAG method to image FRET cleavage (**Fig S3B**). The images demonstrated localization of the fluorescence signal generated by MAPK1 to cytoplasm with sufficient transfection efficiency.

**Figure 3:**
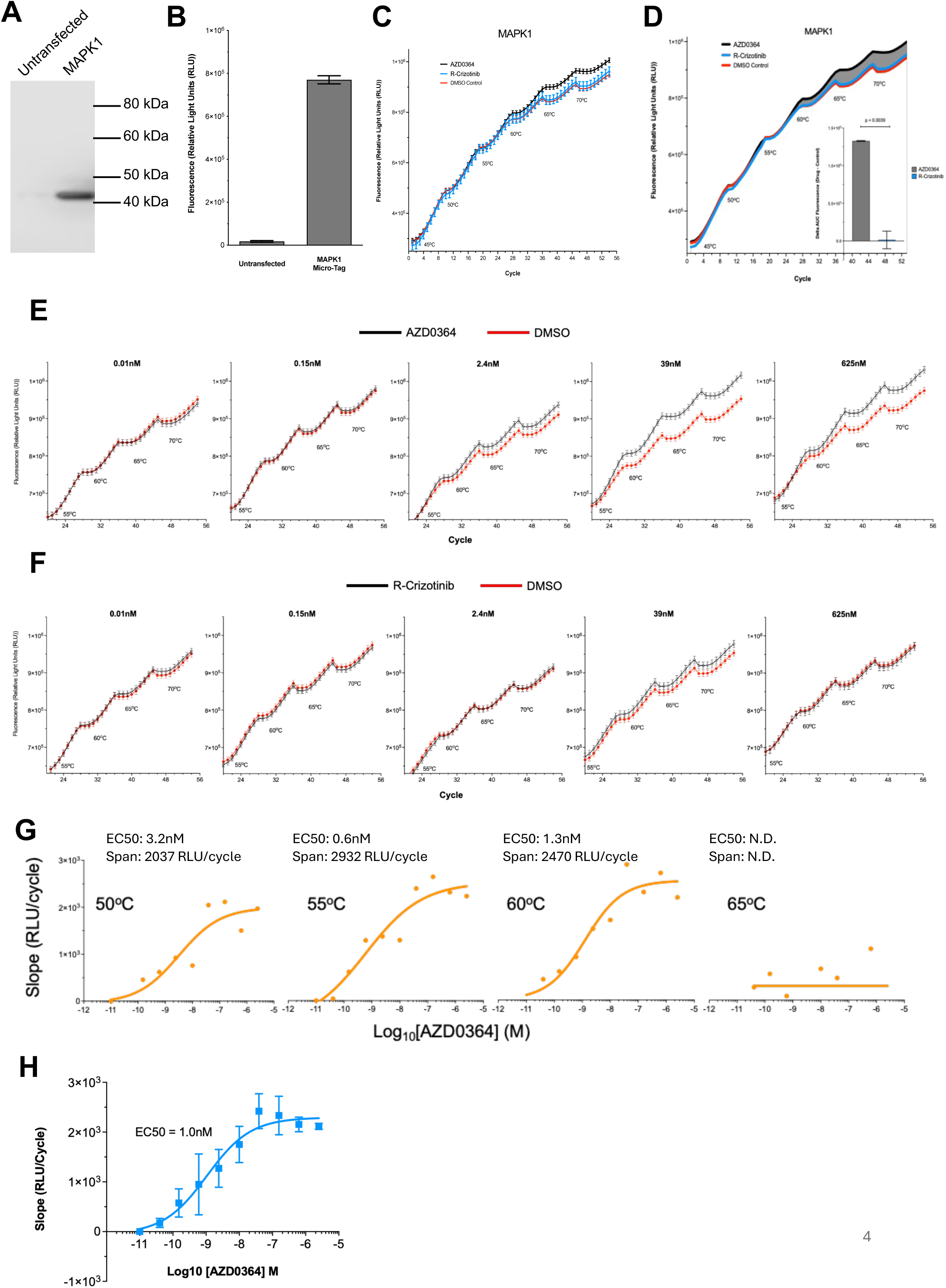
Target Engagement Potency of AZD0364 with MAPK1. (A) Immunoblot of MAPK1-S-tag fusion probed with anti-S-tag antibody. Target is compared to untransfected control. (B) Fluorescence signal measured from enzyme complementation of the target. Target is compared to untransfected control. (C) Temperature series readout from real-time qPCR system measured for MAPK1. Profiles of 1uM AZD0364 (positive control / tool compound), R-Crizotinib (negative control) and DMSO control are shown. (D) Area under the curve (AUC) calculated for the curves in (C) and DMSO AUC subtracted. The inner histogram signified the shaded area for both AZD0364 and R-Crizotinib control. (E) Temperature series profile of increasing doses of AZD0364 tested against various doses of MAPK1. (F) Temperature series profile of increasing doses of R-Crizotinib tested against various doses of MAPK1. (G) Per-temperature EC50 determined from slope of signal generation (RLU/cycle) at each temperature tested. Shown are temperatures 50°C to 65°C. (H) EC50_total_ for MAPK1-AZD0364 engagement from temperatures 50°C to 60°C from 3G above. Experiments were performed in triplicate and error bars represent standard deviation of the mean.

We set to measure cell target engagement of MAPK1 with AZD0364, a commercially available and well-characterized ligand of MAPK1, together with R-Crizotinib (negative control) and DMSO (background control). We hypothesized that such a potent and specific drug-target interaction would yield strong cellular target engagement profile in the MICRO-TAG system. Measurement in temperature series format yielded a unique curve pattern, where all three treatments (at 1uM concentration of AZD0364 and R-Crizotinib) followed an identical curvature up to the 55°C heat step (**Fig 3C** and **3D**). Above 55°C, the curves diverged, indicating that MAPK1 remained soluble due to thermodynamic stabilization imparted by the presence of AZD0364. This separation of signal at 55°C heat step correlates with the determined and reported T_agg_50 of 54°C for MAPK1 (**Fig S3 C and D**) [27]. Area under the curve (AUC) was calculated for both AZD0364 and R-Crizotinib and delta fluorescence above DMSO control was determined by subtracting the AUC for DMSO from treatments (**Fig 3D**). The delta AUC demonstrates significant stabilization of the MAPK1 target by AZD0364 but not the control compound R-Crizotinib (**Fig 3D**).

Dose-response analysis for the treatments demonstrated clear dose-dependent stabilization of MAPK1 by AZD0364, while R-Crizotinib failed to increase the target-driven fluorescence above the DMSO control even at higher doses (**Fig 3E and 3F**). The temperature series protocol allows for determination of per-temperature EC50 values, together with target stabilization levels (**Fig 3G**). The calculated per-temperature EC50 values for MAPK1-AZD0364 engagement were 3.2nM, 0.6nM and 1.3nM for the 50°C, 55°C and 60°C temperature segments, respectively. The span values were 2037 RLU/cycle, 2932 RLU/cycle and 2470 RLU/cycle for these temperature segments. For 65°C segment, values were not determined. While the span stayed consistent, EC50 values fluctuated by 3-fold across these temperatures. The data shows that temperatures between 50°C - 60°C can serve to provide an EC50_total_ value of 1.0nM across multiple segments (**Fig H**). EC50_total_ yields informative insight into the cumulative engagement potency of AZD0364 with MAPK1 across its multiple conformational states. The EC50_total_ value is consistent with the previously reported affinity of AZD0364 for MAPK1 [28]. The conventional T_agg_50-based CTE measurement MAPK1 reporter showed similar results as the temperature series but had less of a range and increased error, overall demonstrating the advantage of the temperature series (**Fig S3 E and F**).

### KRAS case study: Selectivity of MRTX-1133 for KRAS G12D mutant vs wild-type

KRAS represents another type of challenging target, a membrane-bound proto-oncogene that orchestrates complex cancer-driving cellular events. Its relatively smooth protein structure, lack of deep pockets for small-molecule binders and high affinity for GTP add to the challenge. From a therapeutic perspective, high selectivity needs to be achieved for its mutant version as compared to the wild-type (WT) version [29]. We set to generate two parallel MICRO-TAG reporter cells by transient expression of human KRAS (G12D) and KRAS (WT) (Uniprot: G3V5T7) with cloned S-tag (**Fig S4A and S4B**). An initial assessment of expression of the constructs in HEK293 cells resulted in single-band expression of 22 kDa on immunoblot and significant fluorescence signal from enzyme complementation (**Fig 4A and 4B**). Live-cell images of the reporter cells demonstrated that both KRAS (G12D) and KRAS (WT) localized predominantly to the cytoplasm and plasma membrane, albeit at varying intensities consistent with level of expression (**Fig S4C**).

**Figure 4:**
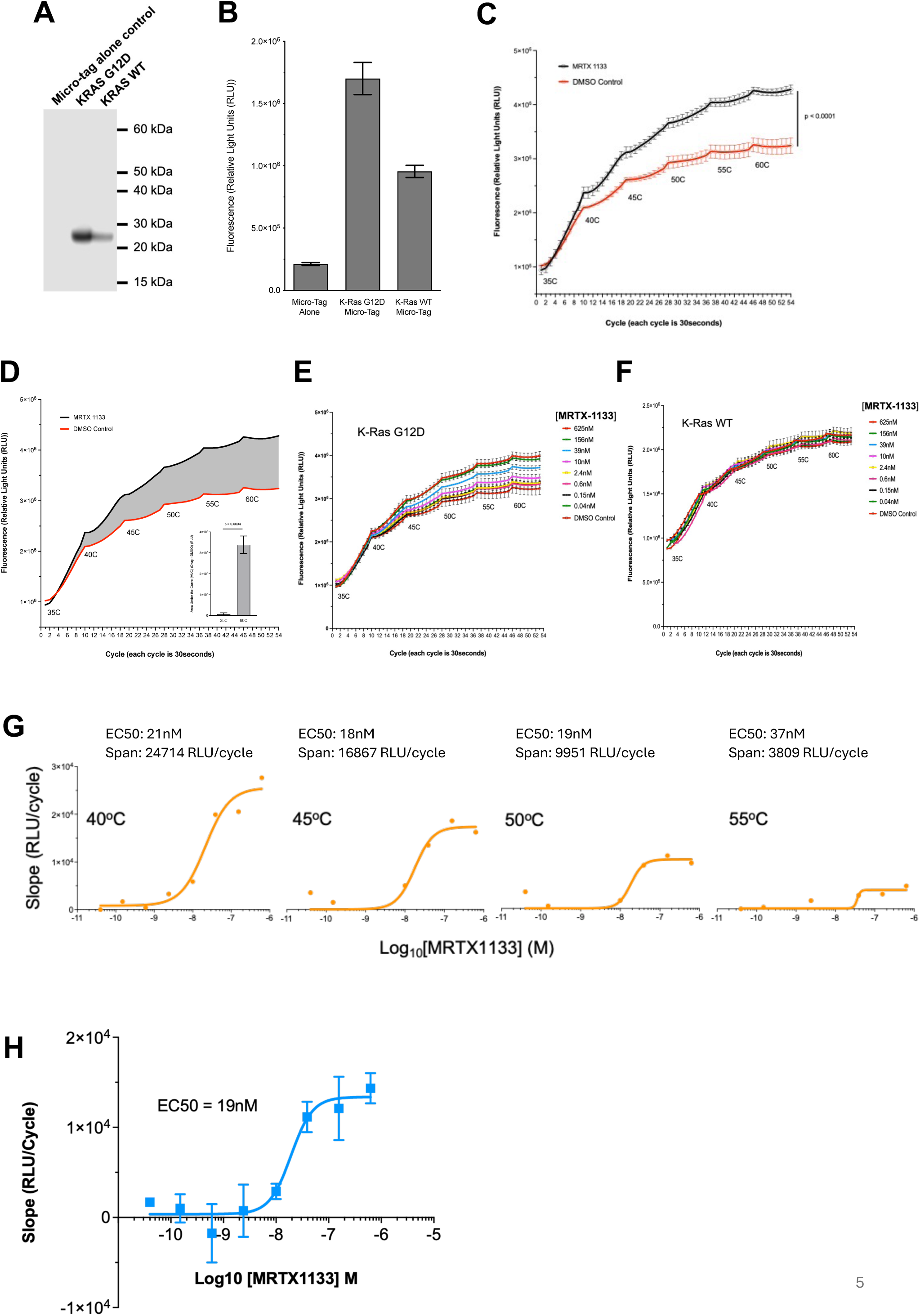
Selectivity of MRTX-1133 for mutant vs wildtype KRAS. (A) Immunoblot of S-tag KRAS (WT) and KRAS (G12D) probed with anti-S-tag antibody. Targets are compared to untransfected control. (B) Fluorescence signal measured from enzyme complementation of the targets. Targets are compared to untransfected control. (C) Temperature series real time curves for KRAS (G12D) with 2.5uM MRTX-1133. Profiles of MRTX1133 (positive control / tool compound) with DMSO control are shown. (D) Area under the curve (AUC) calculated from (D). The inner histogram signified the AUC after 35°C heating step compared with AUC after 65°C heating step. (E) Temperature series profiles of increasing doses of MRTX1133 tested against KRAS (G12D). (F) Temperature series profiles of every tested dose of MRTX1133 tested against KRAS (WT). (G) Per-temperature EC50 determined from slope of signal generation (RLU/cycle) at each temperature tested. Shown are temperatures 40°C to 55°C. (H) EC50_total_ of binding for MRTX1133 to KRAS G12D from temperatures 40°C to 55°C from 4G above. Experiments were performed in triplicate and error bars represent standard deviation of the mean.

We employed MRTX-1133, a characterized noncovalent small molecule inhibitor of mutant KRAS, for cell target engagement to distinguish between the wild-type and mutant forms of KRAS. It is well established that MRTX-1133 binds the (G12D) mutant KRAS with 1000-fold higher selectivity than WT KRAS [30]. The G12D mutation induces distinct conformational changes and altered loop surrounding the binding site in the KRAS protein, creating a highly selective binding pocket for MRTX-1133 [31]. We set to interrogate whether the thermal stability imparted by aromatic interactions between MRTX-1133 and KRAS (G12D) can be captured using the MICRO-TAG system.

Real-time temperature series assay of cell target engagement of MRTX-1133 with KRAS (G12D) and KRAS (WT) yielded expected results. At 2.5uM dose, the inhibitor demonstrated significant stabilization of KRAS (G12D), with a wide signal separation from the DMSO control starting around 40°C (**Fig 4C**). The fluorescence signal continued to increase until 55°C before it plateaued, while signal for the DMSO control plateaued before 50°C. The area between the curves demonstrates the magnitude of target stabilization (**Fig 4D**). Dose-response test of MRTX-1133 against both KRAS (G12D) and KRAS (WT) demonstrated significant selectivity of the inhibitor for the mutant KRAS target (**Fig 4E** and **F**). Deeper analysis of each data segment revealed temperature-dependent engagement profile of KRAS G12D with MRTX1133 (**Fig 4G**). While the per-temperature EC50 values stayed consistent at ∼19nM between 40°C-55°C, the signal span fluctuated between 3809 and 24715 RLU/cycle, demonstrating the thermodynamic shift the target underwent during this temperature series. An EC50_total_ value across all tested temperatures shows binding of MRTX1133 to KRAS G12D of 19nM (**Fig 4H**). EC50_total_ yields valuable insight into the depth of thermodynamic and conformational response of KRAS G12D under proteomic conditions.

### UBE2N case study: Potency of covalent UC-764865 inhibitor relative to structurally related analog

UBE2N presents another challenging target we tested using the MICRO-TAG method. UBE2N, a ubiquitin E2 conjugating enzyme, orchestrates ubiquitination of substrates [32]. UBE2N is involved in various cellular processes, including innate immune and inflammatory signaling, DNA damage response, and mitophagy [32]. In addition, inhibition of UBE2N suppresses growth of several cancers, such as ovarian, breast, and colon cancers, and leukemia and lymphoma [33–35]. Since UBE2N is an emerging target in a variety of cancer and inflammatory diseases, there has been growing interest in developing selective and potent inhibitors. We recently reported a selective small molecule inhibitor of UBE2N that covalently binds to its catalytic site cysteine [20].

Initial assessment of MICRO-TAG reporter cells transiently expressing UBE2N (Uniprot: A0A3P8V155) yielded an expected single-band of 17 kDa by immunoblot (**Fig 5A and S5A**). Enzyme complementation of the target generated significant fluorescence relative to untransfected control (**Fig 5B**). Imaging of live HEK293 cells expressing the tagged UBE2N showed high degree of fluorescence generated by target-driven enzyme complementation (**Fig S5B**). Two compounds, UC-764865 (UC65) and PO1788, were selected for testing against UBE2N due to their high structural similarity (**Fig S5C**). The primary distinction between them lies in UC65’s demonstrated covalent affinity for UBE2N [23]. The MICRO-TAG cell target engagement method was used to compare potency of the two structurally similar compounds for UBEN2. Temperature series and dose response CTE profile of the novel inhibitor UC65 diverged from that of DMSO control at 250nM and plateaued above micromolar doses (**Fig 5C**). The profile of the inactive analog, PO1788, did not generate a meaningful difference compared to the control (**Fig 5D**).

**Figure 5:**
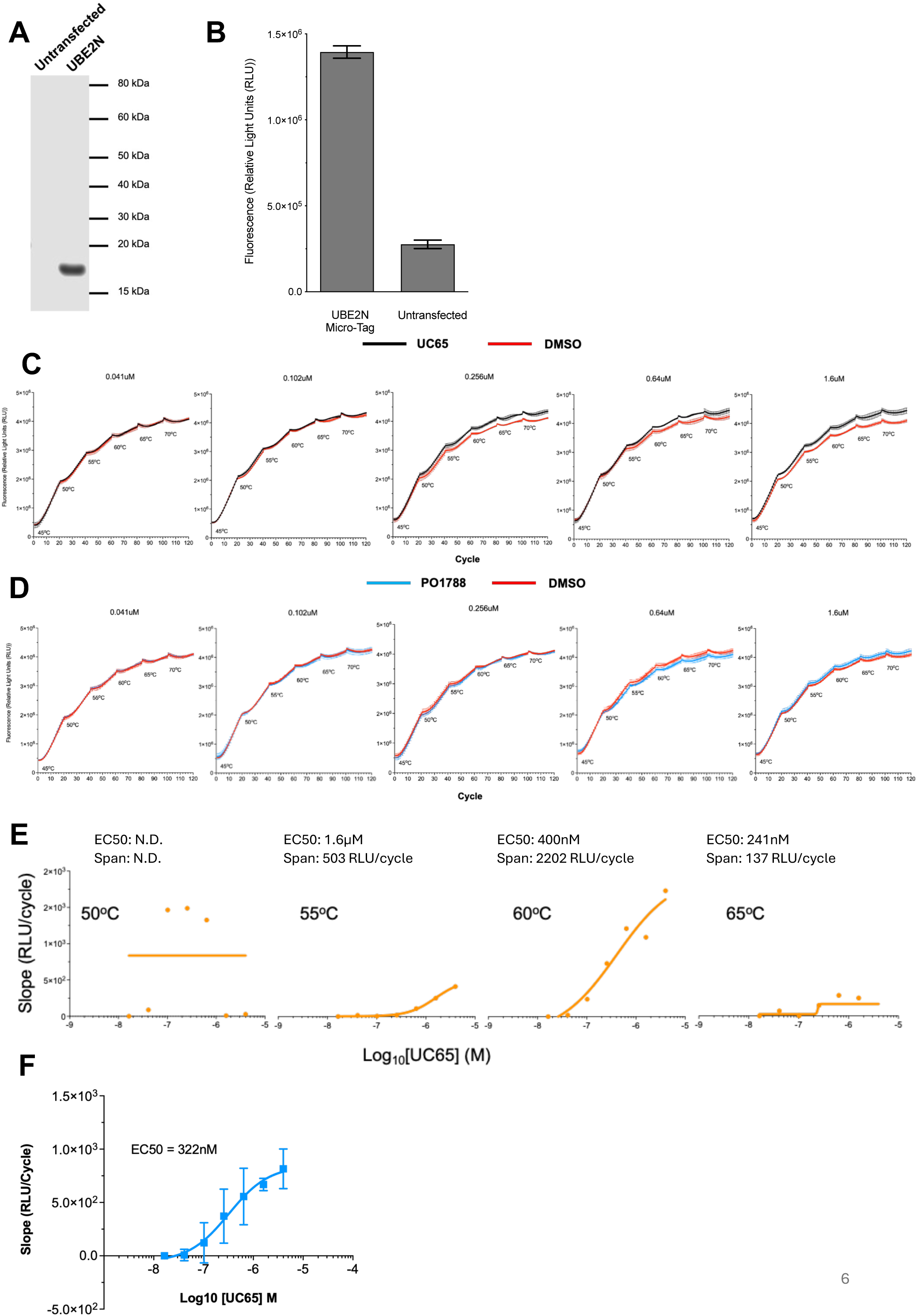
Potency of covalent UC65 and inactive analog with UBE2N. (A) Immunoblot of S-tag UBE2N probed with anti-S-tag antibody. Target is compared to untransfected control. (B) Fluorescence signal measured from enzyme complementation of the target. Target is compared to untransfected control. (C) Temperature series curves from the qPCR real-time system measured for various doses of UC65 compound. (D) Temperature series curves from the qPCR real-time system measured for increasing doses of PO1788 compound. (E) Per-temperature EC50 of binding determined from slope of signal generation (RLU/cycle) at each temperature tested. Shown are temperatures 50°C to 65°C. (F) EC50_total_ of binding for UC65 to UBE2N from temperatures 55°C to 65°C from 5E above. Experiments were performed in triplicate and error bars represent standard deviation of the mean.

The temperature series profile shows UBE2N has a narrow temperature window for determination of target engagement (**Fig 5E**). While the target displayed EC50 of 1.6µM at 55°C, it shifted to 400nM at 60°C, then to 241nM at 65°C. During this course, the signal span fluctuated from 503 RLU/cycle to 2202 RLU/cycle, then down to 137 RLU/cycle at these temperatures, respectively. At 50°C, the target did not display ant significant engagement event. EC50_total_ of 322nM, determined from temperatures 55°C to 65°C, is consistent with what has been reported for UC65 binding to UBE2N (**Fig 5F**) [20]. In comparison to the conventional T_agg_50-based CTE method for UBE2N, the temperature series approach demonstrates a significant increase in signal dynamic range (**Fig S5D-F**). Our data demonstrates the MICRO-TAG temperature series method may be advantageous for increasing sensitivity and dynamic range in CTE experiments for various challenging targets.

## DISCUSSION

Many drug targets exist in the cell in various transient structural and functional states, translating to context-dependent thermodynamic events, which are of high importance for precision therapeutic targeting. The variety in structural variations broadens the spectrum of possible precision drug targeting events, extending beyond the conventional active site pharmacology. This expands the druggable proteome to challenging targets such as protein-protein interactions and allosteric sites, thus offering new avenues for therapeutic intervention. Current CTE methods for assessing drug target interaction in cells, while highly valuable, read end-point signal for drug-target engagement under a single temperature of aggregation (T_agg_50) and hence provide a static and rather simplified view of target profile, averaging the plurality of conformational states of drug targets. Reliance on conventional luminescence-based CTE systems based on single T_agg_50 point carries an inherent bias and misses potential alternative drug-target engagement events in the cell. Real-time detection of fluorescence signal is preferable over end-point signal, particularly due to its higher sensitivity, dynamic range and precision of response quantification [36–38]. The MICRO-TAG system is specifically designed for signal detection with real-time instruments with programmed temperature series format. Every temperature point imparts a gradual structural perturbation onto the target, thus challenging its engagement with the tested drug molecule. Upon incubation at 25 °C after every temperature challenge, drug-target equilibrium is reached, and fluorescence signal is developed via RNase S enzyme complementation. This ensures signal readouts at every temperature point are based on equilibrium at consistent near-physiological temperature. Using Area Under the Curve (AUC) calculation, target engagement response can be quantified enabling calculation of target engagement potency, both per temperature EC50 and EC50_total_. Incubation and signal detection time can be programmatically adjusted to enabling detection of all equilibrium-driven events. However, not all drug engagement events will impart detectable conformational changes on the target of interest, a well-recognized phenomenon for membrane proteins that are thermally protected within membrane structures [39, 40]. In this vein, many promising drug candidates may appear as false negatives. Provided these limitations are addressed, the MICRO-TAG method can be further developed to capture drug-target engagement events at various thermodynamic states, hence enabling measurement of multi-state target engagement events and offering applicability for structurally diverse challenging drug targets.

The fluorescence-based MICRO-TAG enzyme complementation method incorporates the split-RNase S for cell target engagement (CTE). This enzyme is ideally suited for CTE application with real-time systems. Firstly, the enzyme possesses a detachable small subunit, with low hydrophobic and structurally inert content with minimal size of 15 amino acids, which can be used as a tag cloned to a target protein of interest. Second, its large subunit is sufficiently thermally stable to withstand high temperature challenge without losing activity and is catalytically non-functional in the absence of the small subunit. Third, complementation of the small subunit with the large subunit is highly specific and stable enough to enable unbiased measurement of drug-target engagement. RNase S is in a unique position with its utility for fluorescence-based cellular target engagement. Majority of split enzymes cannot be ideal candidates for such an assay because these prerequisites are poorly met [41]. The widely adopted HiBiT system, which is based on NanoLuc, uses a 11-amino acid small subunit predominantly polar and hydrophobic residues, in some cases requiring addition of spacers to extend the tag and neutralize overall charge [42]. Similarly, the β-galactosidase system also employed for CTE uses a 45-amino acid small subunit, which significantly impacts thermal profile of the target with potential misrepresentation of potency profile of tested ligands [43, 44].

The 15-amino acid S-tag cloned to a target protein of interest offers advantages for utility in CTE systems. The unique inert feature of S-tag makes the MICRO-TAG method potentially target-agnostic, enabling its use for a large class of structurally diverse target families that depend on the cellular proteomic context. S-tag has a predominantly hydrophilic and non-polar amino acid composition, which ostensibly allows for the tag to be exposed on the surface of the target protein rather than buried within a hydrophobic pocket [23]. This enables the utility of S-tag with challenging targets such as transmembrane proteins and transcription factors. However, one limitation of a hydrophilic tag for some targets may be narrow signal detection range, as the tag may not completely conceal with an aggregating target, thus still contributing to complementation with its large subunit. To address this limitation, the real-time temperature series method described here can be programmed to introduce empirically tested temperature and incubation conditions to maximize the signal window when testing such challenging targets.

The MICRO-TAG method is based on an enzymatic reaction for fluorescent signal development, detected by highly sensitive real-time instruments. This provides high signal-to-noise ratio together with improved sensitivity and dynamic range, as compared to other similar strategies. An enzymatic reaction generating fluorescent signal amplifies readout allowing for detection of lower levels of target protein, thus resulting in a higher dynamic range. In addition, versatility is incorporated into the assay with choice of various donor-quencher pairs for the FRET substrate. The MICRO-TAG method is amenable to various wavelengths for detection, thus allowing for multiplexing capability with other florescence-based techniques and enhancing cellular target engagement readouts. Fluorescence offers advantages over luminescence, particularly in terms of its high sensitivity, selectivity, and faster response time, as fluorescence requires an external light source to excite molecules, allowing for precise control over the emission signal and minimizing background noise. This is in contrary with the broader luminescence phenomenon where light emission can occur from various sources without specific excitation [45–48]. Fluorescence can detect very low levels of a target molecule due to the ability to precisely control the excitation light and filter out background noise. By selecting specific donor-quencher pairs with distinct excitation and emission wavelengths, fluorescence allows for selective detection of target molecules within a complex cellular milieu. Fluorescence emission occurs almost instantaneously upon excitation, enabling rapid detection and real-time imaging. Incorporation of fluorescence with CTE allows for expansion to microscopy, flow cytometry, multiplexing assays, and diagnostics due to versatility of the FRET substrate and high sensitivity [12, 49].

The “mix-and-detect” feature of the MICRO-TAG system obviates the requirement for pre-incubation, enables faster turnaround time and provides EC50_total_ readouts of drug-target engagement. “Mix-and-detect” systems have been widely adopted for discovery, diagnostic and clinical applications, mainly thanks to their simplicity, cost effectiveness and scalability [50–52]. When integrated with readily available real-time instruments, these advantages will reduce the per-compound cost during early drug discovery screening, accelerate drug discovery efforts at scale, and thus enable efficient identification of promising therapeutic candidates.

Advantages and disadvantages of the MICRO-TAG system warrant further research. Though the scope of this study was limited to introduction and utility of this method in the context of real-time temperature series CTE, it can be potentially expanded for more advanced real-time applications. The currently limited cell penetration quality of the RNA FRET substrate will need to be optimized to enable future specialized applications. To that end, various transfection conditions and substrate modifications are currently being explored to improve its stability and cellular availability. Such advancements of the MICRO-TAG system will enhance various applications ranging from gauging protein-protein interactions to specific therapeutic target requirements [53–60]. The future versions of the MICRO-TAG system will also address the current desire to monitor drug-target engagement to capture disease-relevant proteomic events in real time, with direct utility for various stages of drug discovery [61–64]. Further applications may include live-cell imaging for deep insight into therapeutic mechanism-of action, multiplexing of various fluorescent signals for enhanced visualization of intricate drug-target interactions and complexes, together with ability to study other dynamic drug-target engagement events.

## CONCLUSIONS

MICRO-TAG system, based on split-RNAse S enzyme complementation, offers a novel fluorescence-based CTE method for early drug discovery. It enables sensitive real-time measurement of drug-target engagement within the cellular context. By virtue of a programmable temperature series method, it bypasses the need to empirically establish the thermal melting profile of the target protein prior CTE measurements, thus also removing the T_agg_ bias. This procedurally scalable method is particularly suited for structurally diverse targets without reliance on predetermined T_agg_50 point. The MICRO-TAG system empowers assay setup and analysis through increased sensitivity and flexibility of real-time instrument programming. This methodology streamlines drug discovery workflows offering savings on time and cost and allowing for improved throughput and ease of automation while enabling discovery for therapeutics to challenging drug targets.

## METHODS

### Cell lines

HEK293T (ATCC CRL-3216) cells were cultured in DMEM (Gibco) supplemented with 10% fetal bovine serum (FBS, Gibco) and 1% (v/v) Penicillin-Streptomycin (Gibco) in a 37C humidified incubator with 5% CO2. The cells were passaged every 2–3 days when they reached approximately 80%–90% confluence. Transfection of cells was by reverse transfection with LF3000 (Thermo Fisher Scientific) following manufacture protocol for 6 well plate transfection. 48hrs post-transfection cells were used for cell target engagement study.

### Chemicals

AZD0364, R-Crizotinib, S Crizotinib, and MRTX-1133 were purchased from Selleck Chemicals. The UBE2N inhibitors UC-764865 and negative control (PO1788) were previously described (35263148, 37102608). UC-764865 and PO1788 were synthesized at Wuxi AppTec. Chemical structure of the compound was analyzed by nuclear magnetic resonance (NMR).

### Target expression constructs

Target proteins human MAPK1 (Uniprot: P28482), human KRAS G12D and WT (Uniprot: G3V5T7) and UBE2N (Uniprot: A0A3P8V155), were cloned into pcDNA1 vector with S-tag either on N- or C-terminus (**Fig S3-5**). Constructs were transfected into HEK293 cells using Lipofectamine 3000 for 48 hours. Subsequently, cells were tested for target expression and used for MICRO-TAG enzyme complementation.

### Synthesis of S-protein

S-protein was cloned into pET28A with 6xHIS tag on C-terminus and expressed in BL21 (DE3) *E. coli* with 0.1 mM IPTG (Isopropyl β-D-1-thiogalactopyranoside) induction. Soluble fraction was separated and expressed S-protein was purified using Ni-NTA affinity purification method. S-protein was eluted using imidazole gradient. Purified protein was checked for purity using Coomassie staining (**Fig S1**).

### Synthesis of FRET substrate

FRET RNA substrate was the optimized fluorogenic substrate 6-FAM-dArUdAdA-6-TAMRA, where 6-FAM (6-carboxyfluorescein) and 6-TAMRA (6-carboxy-tetramethylrhodamine) were added to ribonucleotides by Integrated DNA Technologies (IDT).

### Non-denaturing cell lysis and immunoblot analysis

Cells expressing the S-tag construct were grown in a 6 well tissue culture plate. The cells were lifted by pipetting up/down then transferred to 15ml tube. Cells were pelleted at 400g for 3 minutes, washed with TBS (Tris Buffered Saline (1X TBS; 150mM NaCl, 50mM Tris-HCl pH 7.4) (Thermo Fisher Scientific; cat J62938.k7) and pelleted again. TBS wash removed and cells lysed with 300ul non-denaturing lysis buffer (non-denaturing Lysis Buffer: 1% Triton X-100 (in TBS) containing cOmplete, Mini, EDTA-free Protease Inhibitor Cocktail (Roche; cat #11836153001) for 1 hour at 4°C on a rotator. The lysate was clarified by centrifugation at 14,000rpm for 1minute in a micro-centrifuge. Lysates were used for immunoblot analysis with primary antibody S-Tag (D2K2V) XP Rabbit mAb #12274 from Cell Signaling Technology.

### MICRO-TAG Enzyme Complementation: Temperature series CTE assay using non-denaturing lysates

Test ligands were prepared in DMSO at 100X. Non-denaturing lysates were diluted 1/20 with cold TBS on ice and aliquoted 49.5ul/well to 12 wells of a 96-well PCR plate and 0.5ul of the 100X diluted ligands was added. The 60ul of reaction buffer containing S-protein and FRET RNA substrate were added to a MicroAMP Optical 96-well reaction plate and 20ul of lysates + ligand added. FAM fluorescence (excitation 493nm; emission 517nm) detected with programmed temperature series using Applied Biosystems QuantStudio 3 Real-Time System (Thermo Fisher Scientific). Experiments were performed in triplicate.

### MICRO-TAG Enzyme Complementation: Temperature series CTE assay using intact cells

Transiently transfected HEK293 cells 48hrs post-transfection grown in a 6-well plate were lifted by pipetting up/down and transferred to a 15ml tube. Cells were pelleted 400g 3min centrifugation and washed with TBS and pelleted again. Cells were resuspended in 1ml TBS and aliquoted 49.5ul/well to 12 wells of a 96-well PCR plate and 0.5ul of the 100X ligands diluted in DMSO was added and incubated on ice for 1 hr. For detection, 50ul of 0.5% Triton-X-100 was added to the cells and 20ul of this added to 60ul of reaction buffer containing S-protein and FRET RNA substrate in wells of a MicroAMP Optical 96-well reaction plate. FAM fluorescence (excitation 493nm; emission 517nm) was detected with the temperature series programming using Applied Biosystems QuantStudio 3 Real-Time System (Thermo Fisher Scientific). Experiments were performed in triplicate.

### Live-cell imaging

24hrs post transfection of HEK293 cells with the S-tag protein expression construct, the cells were lifted from the wells of a 6 well plate with trypsin and transferred to 15ml tube. Cells were pelleted by centrifugation at 400g 3min, supernatant removed, and cells resuspended in 3ml of media. Cells were distributed (500ul/well) of a 12 well plate. 24hrs later the media was removed and replaced with 400ul Opti-MEM. For each well, 900pmoles FRET RNA was added to 50ul of Opti-MEM. LF3000 transfection reagent (3ul) is added to 50ul of Opti-MEM; then combined with the diluted FRET RNA and incubated at room temperature for 10min then add dropwise to the cells. After 20min incubation at 37°C the cells were imaged with EVOS M5000 imaging system. FAM fluorescence (excitation 493nm; emission 517nm).

### Quantification and statistical analysis

All experiments were performed in triplicate and are presented as mean ± standard deviation (SD). Statistical analyses were conducted in GraphPad Prism v10.1.1. Group differences were evaluated by two-way ANOVA followed by post-hoc t-tests, with statistical significance defined as p < 0.05. Error bars denote SD of triplicates. For quantitative summaries, both Area Under Curve (AUC)-based and slope-based approaches were applied to each temperature segment (or to entire temperature series), then compared across drug concentrations.

#### AUC-based quantification

For each drug concentration x and temperature segment k with monotone time points (t_i, y_i), area under the curve (AUC) was computed by the trapezoidal rule:

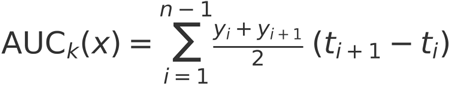

Optional baseline subtraction was applied before integration when specified:

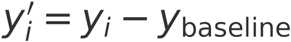

Drug effect relative to DMSO control was quantified as delta AUC per temperature segment:

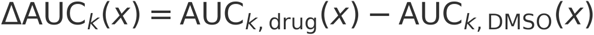

To summarize across temperatures, per-segment AUC values were aggregated using predefined weights w_k (default equal weights):

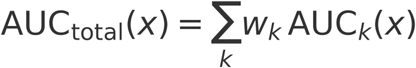

Weights constraint:

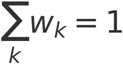

#### Slope-based calculation

Within each temperature segment k, rate of signal generation at drug concentration x was estimated by ordinary least squares (OLS):

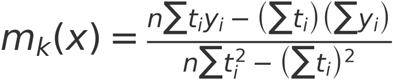

Control-corrected rate when used:

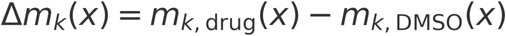

An overall rate across temperatures was computed via weighted aggregation:

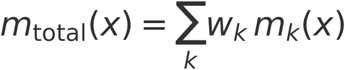

Weights constraint:

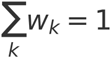

#### Dose–response modeling and EC50 definitions

For both AUC- and slope-derived readouts (raw or delta vs. DMSO), concentration–response curves were fit using the sigmoidal dose–response (variable slope; 4-parameter logistic, 4PL). Per-temperature EC50 values, EC50(T_k), were obtained by fitting a curve at each temperature k:

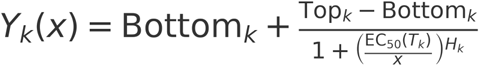

A total potency across the temperature series, EC50_total, was obtained by first aggregating the response across temperatures at each x (e.g., AUC_total(x) or m_total(x)) and then fitting the same 4PL:

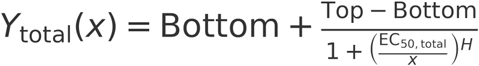

Here, Y(x) is AUC, ΔAUC, m_k, or Δm_k; Bottom and Top are the asymptotes; H is the Hill slope; and EC50 is the concentration producing half-maximal effect.

The aggregate-response approach does not depend on per-temperature EC50 values. Temperature segments where EC50(T_k) is not determined (N.D.) are still included, provided AUC_k(x) or m_k(x) data are available. When some temperatures lack data at specific drug concentrations, weights were renormalized at that concentration so that the aggregated response remains comparable:

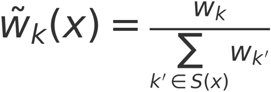

Per-concentration aggregation using the normalized weights (R_k denotes AUC_k or m_k):

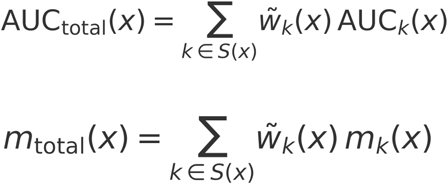

If an entire temperature segment lacked measurable data, w_k was set to 0 and this exclusion was noted. Sensitivity analyses compared EC50_total_ from the full dataset (with per-x renormalization) to EC50_total_ computed on the subset of concentrations common to all temperatures.

## Supporting Information

**Table S1:** Biophysical properties of S-tag.

| **Property name** | **Value** |
| --- | --- |
| Tag length | 15 amino acids |
| Tag sequence | KETAAAKFERQHMDS |
| Theoretical pI | 7.7 |
| Net charge | 0 |
| Aliphaticity index | 20.00 (low thermostability) |
| Hydropathicity score | -1.393 (hydrophilic) |

**Figure S1:**
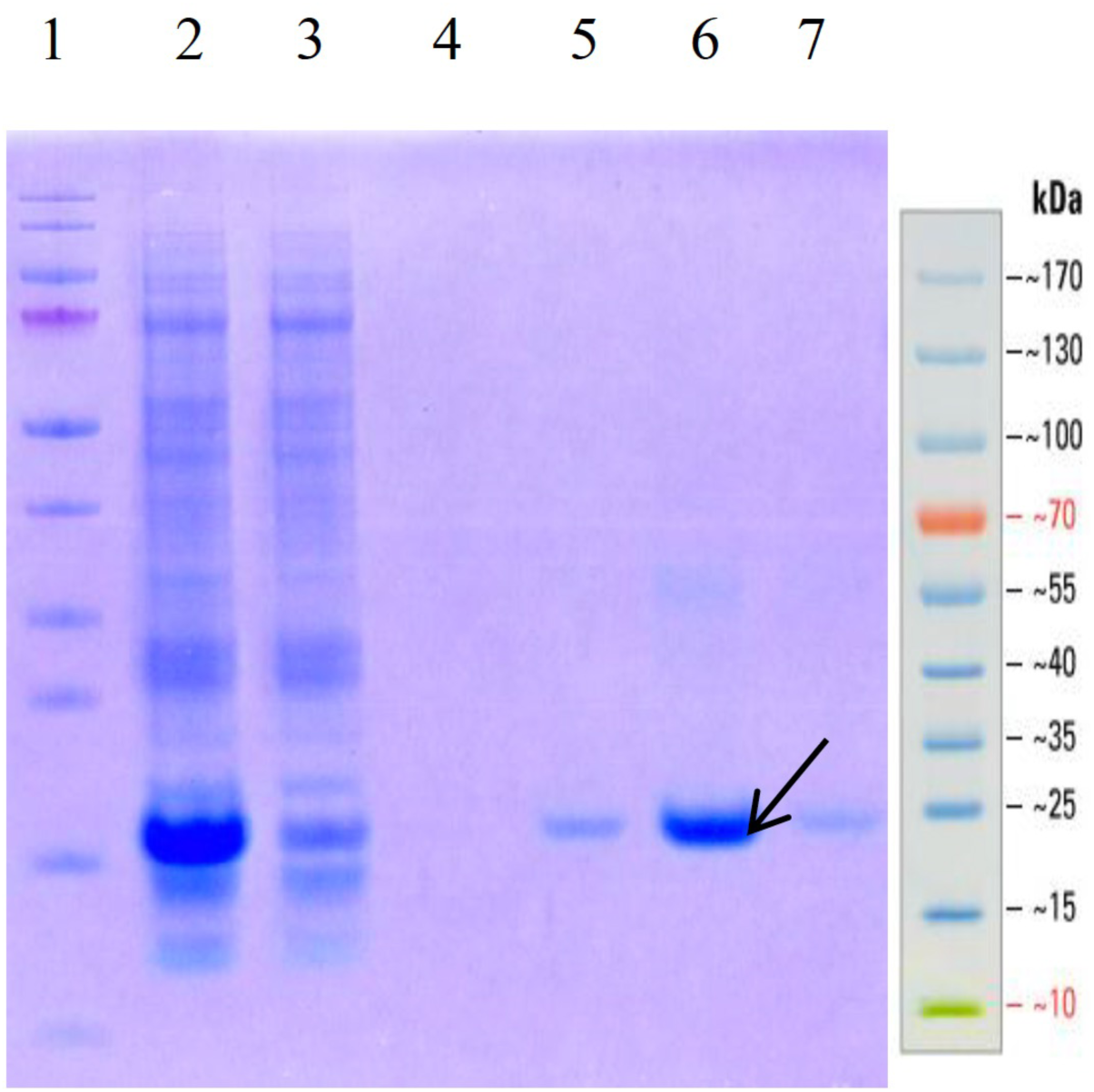
Coomassie staining for purification of recombinant S-Protein.

**Figure S2:**
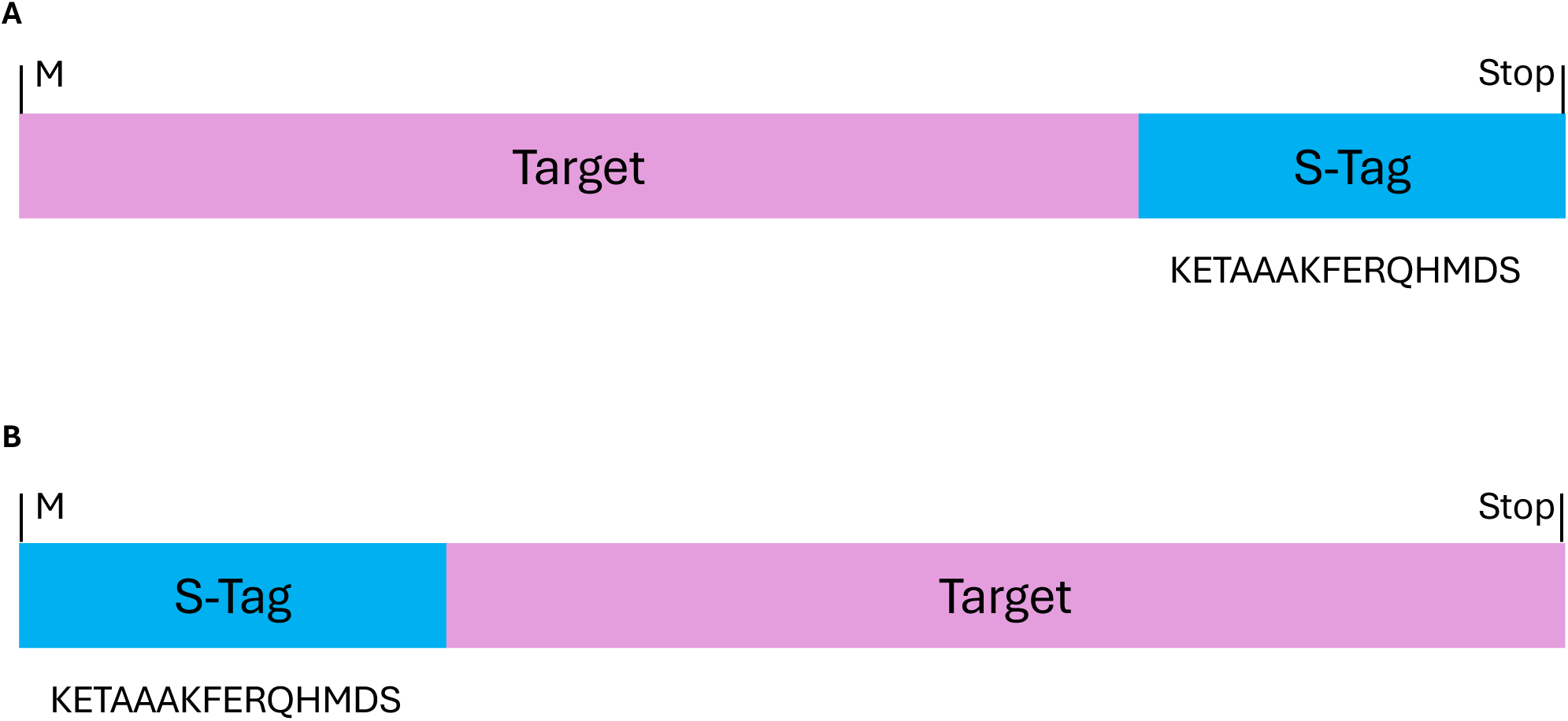
Map of expression construct of MICRO-TAG targets.

**Figure S3:**
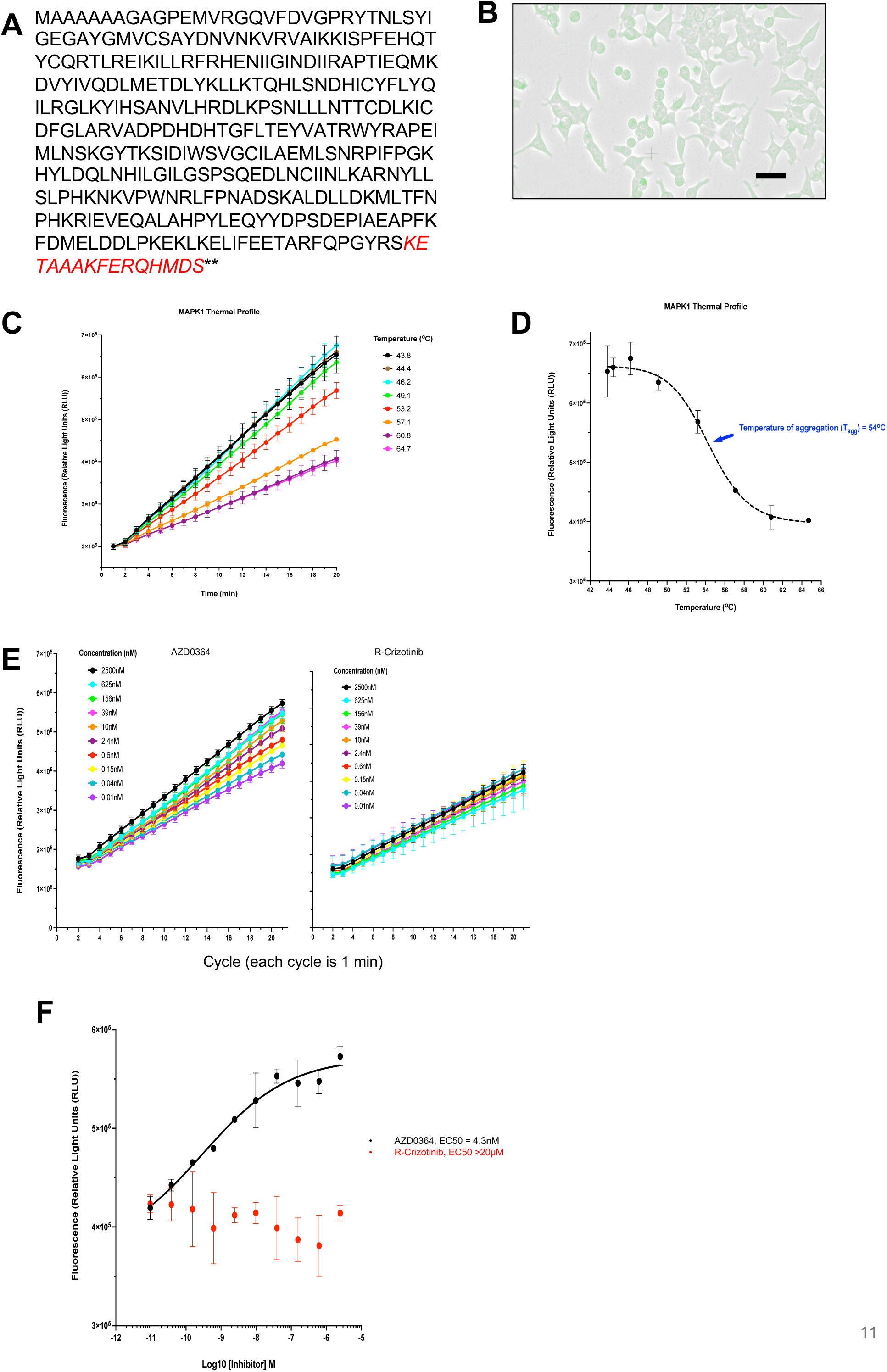
Quality control of cells expressing MICRO-TAG MAPK1 and conventional CTE method for determining EC50 of binding.

**Figure S4:**
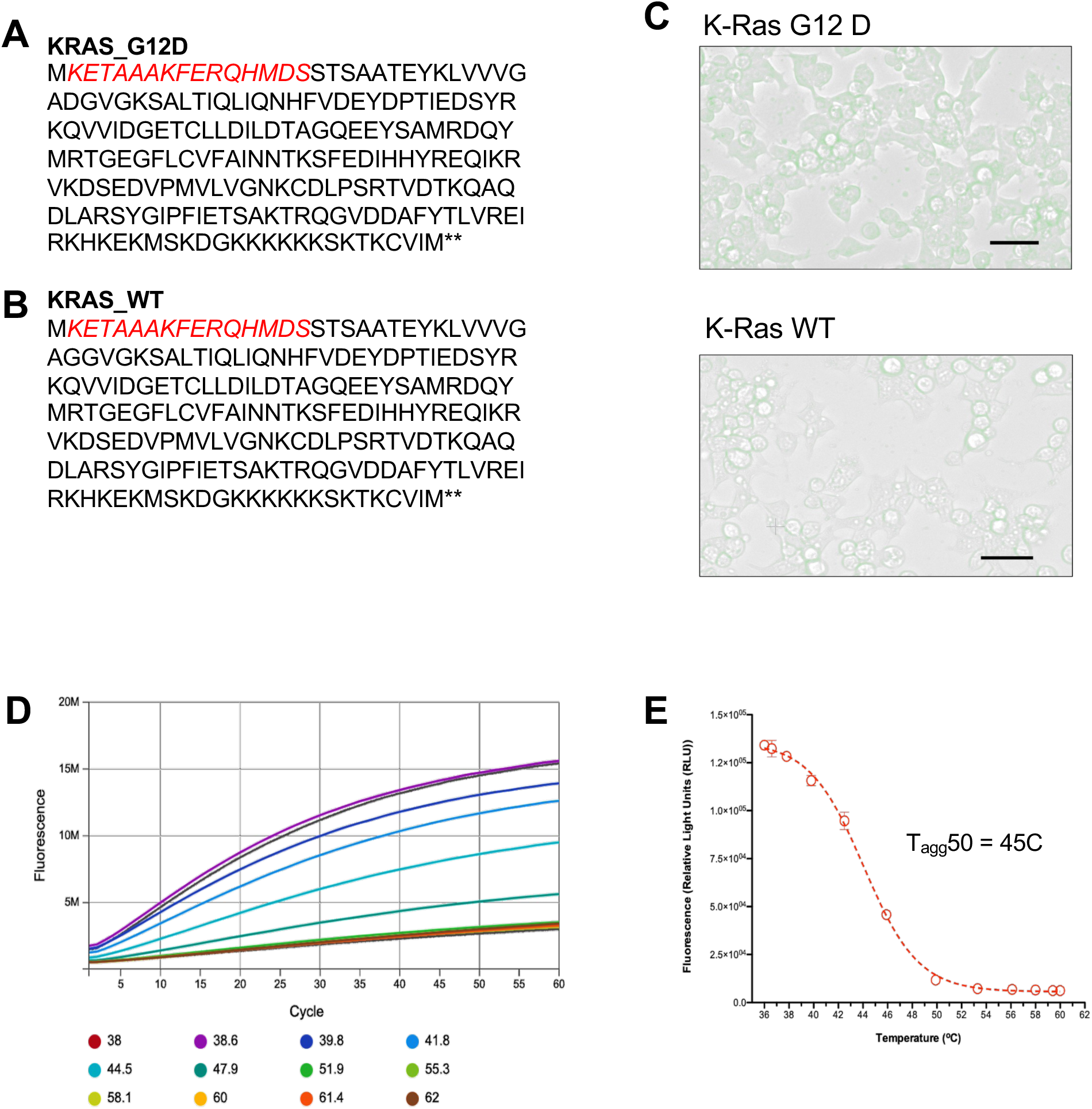
Quality control of cells expressing MICRO-TAG KRAS and conventional CTE method for determining EC50 of binding.

**Figure S5:**
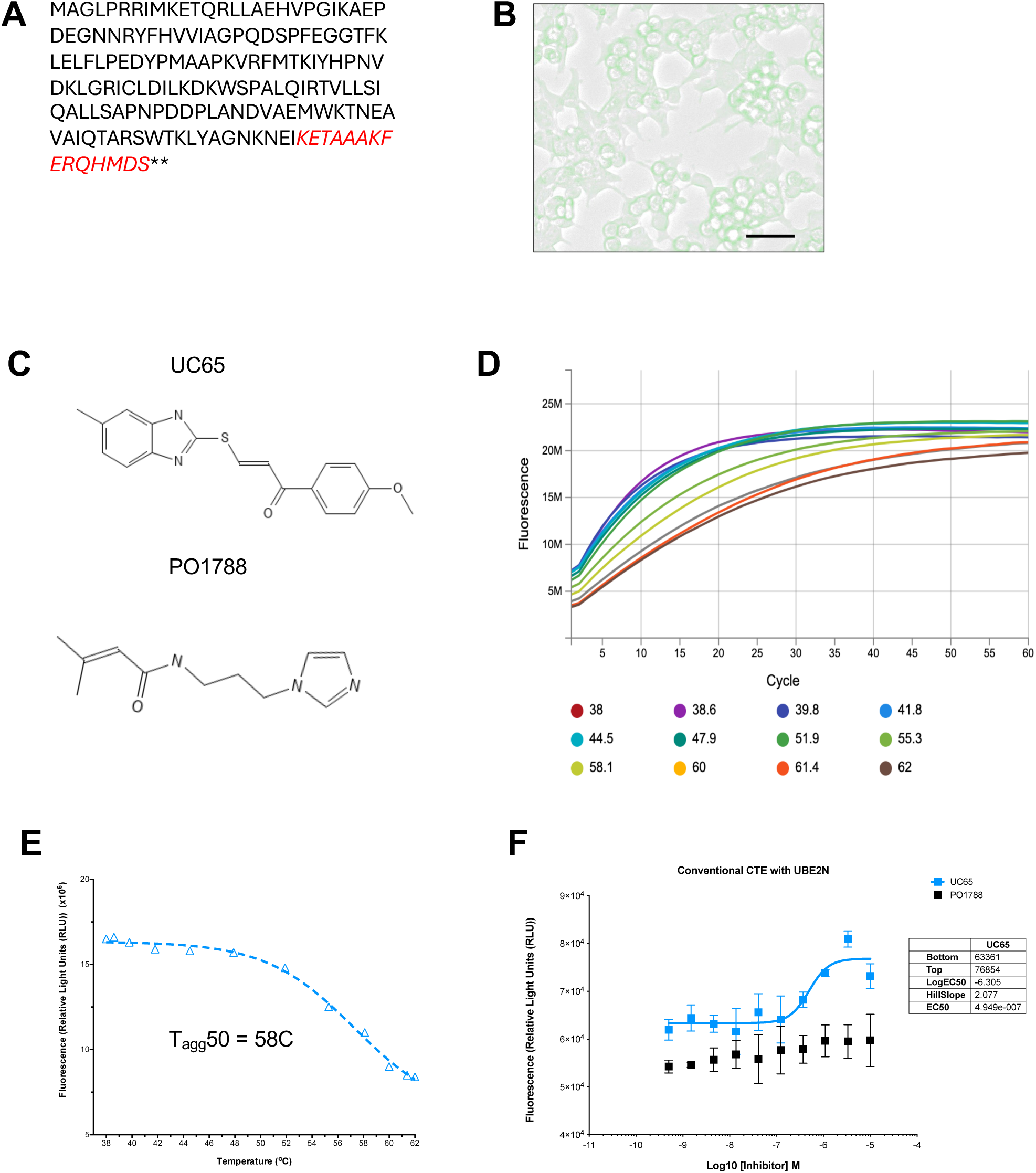
Quality control of cells expressing MICRO-TAG UBE2N and conventional CTE method for determining EC50 of binding.

