## Supporting information for "MICRO-TAG enzyme complementation enables quantification of cellular drug-target engagement in temperature series"

### SUPPLEMENTARY FIGURE LEGENDS

**Table S1: Biophysical properties of S-tag**

Various biophysical parameters of S-tag are provided, particularly, peptide length, peptide sequence, theoretical pI, Net charge, aliphaticity index and hydropathicity score.

**Figure S1: Coomassie staining for purification of recombinant S-Protein**

Lane 1: Marker; Lane 2: Supernatant; Lane 3: Flow through; Lane 4: Elution by 2mM imidazole; Lane 5: Elution by 20mM imidazole; Lane 6: Elution by 250mM imidazole; Lane 7: Beads.

**Figure S2: Expression construct of MICRO-TAG target**

(A) Diagram showing cloning of target with S-tag at C-terminus.

(B) Diagram showing cloning of target with S-tag at N-terminus.

**Figure S3: Quality control of cells expressing MICRO-TAG MAPK1**

(A) Amino acid sequence of MAPK1 cloned with S-tag at C-terminus (shown in red).

(B) Real-time live-cell imaging of enzyme complementation of S-tag MAPK1 with S protein and FRET RNA cleavage in transfected HEK293 cells. Green represents areas with fluorescence of target-induced enzyme complementation.

(C) Real-time MICRO-TAG enzyme complementation data showing thermal melting of MICRO-TAG MAPK1 at various temperatures.

(D) Thermal melting profile of MICRO-TAG MAPK1 derived from (B), showing $T_{agg}50$ of 54$^{o}$C.

(E) Conventional cell target engagement method using Micro-Tag enzyme complementation MAPK1 with AZD0364 (left) and R-Crizotinib (left) at various doses, at 54$^{o}$C.

(F) Cell target engagement potency curves for AZD0364 and R-Crizotinib, extracted from (E), showing EC50 values of 4.3nM and >20µM for AZD0364 and R-Crizotinib, respectively.

**Figure S4: Quality control of cells expressing MICRO-TAG KRAS**

(A) Amino acid sequence of KRAS G12D cloned with S-tag at N-terminus (shown in red).

(B) Amino acid sequence of KRAS wild-type cloned with S-tag at N-terminus (shown in red).

(C) Real-time live-cell imaging of S-tag KRAS (WT) and KRAS (G12D). Green represents areas with fluorescence of target-induced enzyme complementation.

(D) Representative real-time MICRO-TAG enzyme complementation data showing thermal melting of MICRO-TAG KRAS G12D at various temperatures.

(E) Thermal melting profile of MICRO-TAG KRAS G12D derived from (C), showing $T_{agg}50$ of 45°C.

**Figure S5: Quality control of cells expressing MICRO-TAG UBE2N**

(A) Amino acid sequence of UBE2N cloned with S-tag at C-terminus (shown in red).

(B) Real-time live-cell imaging of S-tag UBE2N expressing HEK293 cells. Green represents areas with fluorescence of target-induced enzyme complementation.

(C) Chemical structures of UC65 (active compound) and PO1788 (inactive compound).

(D) Real-time MICRO-TAG enzyme complementation data showing thermal melting of MICRO-TAG UBE2N at various temperatures.

(E) Thermal melting profile of MICRO-TAG UBE2N derived from (C). $T_{agg}50$ of 58°C is shown.

(F) Conventional cell target engagement method using Micro-Tag enzyme complementation for UBE2N with UC65 or PO1788 at various doses, at 58°C. Shown are cell target engagement potency curves showing EC50 values of 495nM for UC65 and no curve fitting for PO1788.
